# Evaluation of a polymeric composite bone filler scaffold for local antibiotic delivery to prevent *Staphylococcus aureus* infection in a contaminated bone defect

**DOI:** 10.1101/2020.12.21.423753

**Authors:** Karen E. Beenken, Mara J. Campbell, Aura M. Ramirez, Karrar Alghazali, Christopher M. Walker, Bailey Jackson, Christopher Griffin, William King, Shawn E. Bourdo, Rebecca Rifkin, Silke Hecht, Daniel G. Meeker, David E. Anderson, Alexandru S. Biris, Mark S. Smeltzer

**Affiliations:** Department of Microbiology and Immunology, University of Arkansas for Medical Sciences, Little Rock, AR; Center for Integrative Nanotechnology Sciences, University of Arkansas at Little Rock, Little Rock, AR; Department of Large Animal Clinical Sciences, University of Tennessee College of Veterinary Medicine, Knoxville, TN; Department of Small Animal Clinical Sciences, University of Tennessee College of Veterinary Medicine, Knoxville, TN

## Abstract

We previously reported the development of an osteogenic bone filler scaffold consisting of degradable polyurethane (dPU), nano-sized hydroxyapatite (nHA), and decellularized bovine bone particles (DBP). In this report we describe the results of studies aimed at evaluating the use of this scaffold as a means of local antibiotic delivery for the prevention of infection in a segmental bone defect contaminated with *Staphylococcus aureus*. We evaluated two different scaffold formulations that contained the same components in the same ratios but differed from each other with respect to overall porosity and therefore surface area. Studies done with vancomycin, daptomycin, and gentamicin confirmed that antibiotic uptake was concentration dependent and that increased porosity was correlated with increased uptake and prolonged release of all three antibiotics. Vancomycin could be passively loaded into either scaffold formulation in an amount sufficient to prevent infection, as evidenced by the complete eradication of viable bacteria from the surgical site of most animals in a rabbit model of a contaminated mid-radial segmental bone defect. Even in those few cases in which complete eradication was not achieved, the number of viable bacteria present in the bone was significantly reduced comparison to untreated controls. There was also no radiographic evidence of osteomyelitis in any rabbit treated with vancomycin-loaded scaffold. Microcomputed tomography (μCT) of bone defects up to 84 days of exposure to scaffolds with and without vancomycin also demonstrated that the addition of vancomycin even in the highest concentration did not significantly diminish the osteogenic properties of either scaffold formulation. Together, these results demonstrate the potential utility of our bone regeneration scaffold for local antibiotic delivery.

## Introduction

Open fractures and penetrating wounds that damage the bone are extremely complex injuries. The resulting bone defects are particularly problematic because they involve a breach of the skin and occur in non-sterile environments, thus making them highly susceptible to infection. The treatment of these injuries requires an intensive interdisciplinary clinical approach that includes long-term systemic antibiotic therapy and surgical debridement to remove damaged and contaminated bone and soft tissues [1–5]. Debridement can exacerbate the bone injury even further, thus increasing the likelihood that reconstruction will be required. This often necessitates the use of indwelling hardware and bone fillers including autografts, allografts, or various forms of synthetic bone void fillers [6]. The compromised wound environment also often makes it necessary to augment systemic therapy using some form of local antibiotic delivery, the goal being to obtain antibiotic levels at the site of injury that exceed those that can be achieved using systemic methods and are high enough to overcome the intrinsic resistance associated with formation of a bacterial biofilm on damaged bone and soft tissues [7–13]. However, despite such intensive medical and surgical intervention, recurrent infection and even amputation are far-too-common outcomes.

Thus, the two most critical clinical concerns in the treatment of traumatic bone injuries are restoring the structural integrity of damaged bone and preventing infection from developing in a contaminated bone defect. Because it is imperative to limit the possibility of recurrent infection following surgical debridement, these critical considerations are also of importance in other forms of osteomyelitis (OM), including those that are initiated by hematogenous seeding of the bone or invasion of the bone from an overlying soft tissue infection [2, 4, 5, 14]. Recent reports have demonstrated that a novel polyurethane-based bone filler scaffold promotes bone regeneration even in critical-sized bone defects and that the scaffold can potentially be used to absorb and release bioactive agents including antibiotics [15–18]. However, to date, its use as an antibiotic-delivery vehicle to prevent infection in a contaminated bone defect has not been experimentally tested in an appropriate animal model.

The scaffold provides a highly flexible platform that allows for precise control of structural and physical characteristics including elasticity, compressibility, porosity and surface area [15–18]. The ability to control porosity is important to help ensure that the antibiotic concentrations achieved are sufficient to prevent infection. However, even local antibiotic delivery can increase systemic antibiotic levels. Consequently, the combined use of systemic and local antibiotic therapy, while clinically necessary with respect to preventing and/or eradicating infection, can have adverse physiological consequences on the host [19, 20]. This makes it important to determine the minimum amount of antibiotic required to reliably achieve the desired clinical result while minimizing the risk of such adverse effects.

To address these objectives, we carried out *in vitro* and *in vivo* experiments to assess two issues. First, we determined whether enough antibiotic can be incorporated into the scaffold to prevent infection in a contaminated bone defect without compromising bone regeneration. Next, we determined the minimum amount of antibiotic required to achieve therapeutic concentrations without compromising bone regeneration.

## Materials and methods

### Scaffold construction

These scaffold variations were prepared by slightly modifying the original method such that the resulting scaffolds (S1 and K20) were built with two highly different BET (Brunauer, Emmet, and Teller) specific surface area values [16]. For K20 the increased surface area was achieved by using a salt leaching approach. In anticipation of *in vivo* studies both scaffolds were produced in a 1 cm X 4 mm architecture to match the surgical defect created in our rabbit model. The surface areas determinations were done by using a BET method and N_2_ adsorption. The instrument used was a Micromeritics ASAP2020 and the measured BET surface area values for S1 was 36.4 m^2^/g and for K20 was 54.7 m^2^/g, respectively.

### Antibiotic loading and elution

To assess the antibiotic loading capacity of each scaffold formulation, individual S1 and K20 scaffolds were chosen at random and soaked at room temperature (RT) in 2.0 mL of phosphate-buffered saline (PBS) containing 1.0, 10.0 or 100.0 mg/mL of vancomycin, daptomycin, or gentamicin, all of which were obtained from the inpatient pharmacy at the University of Arkansas for Medical Sciences. Three scaffolds of each formulation were soaked in a solution containing an antibiotic at the concentrations above. Initially, each scaffold was soaked for 24 hours, although this time period was reduced to 1 hour in later experiments as detailed below. After soaking, the scaffolds were removed from the antibiotic solution and transferred individually to the wells of a 24-well microtiter plate containing 1.0 mL of antibiotic-free PBS. Plates were incubated at 37°C on an orbital shaker with continuous mixing. After 30 min, the buffer was removed and replaced with fresh antibiotic-free PBS. This was repeated every 24 hours thereafter for up to 10 days.

### Antibiotic activity assays

As samples were collected the approximate amount of active antibiotic was determined using a broth dilution assay in which two-fold dilutions of each sample were compared to a standard prepared with fresh antibiotic [12]. This allowed us to determine a two-fold range of active antibiotic in each sample; the results of these bioassays are reported based on the lower end of this two-fold range. Our vancomycin bioassays were verified using a spectrophotometric assay based on absorbance at 280 nm as previously described [25].

### *In vivo* rabbit osteomyelitis model

Our initial *in vivo* studies were done with the K20 scaffold formulation saturated in a solution containing 100 mg/mL of vancomycin based on the results of *in vitro* elution studies demonstrating maximum antibiotic uptake and elution at this concentration (see below). To test *in vivo* relevance, we used a rabbit model of post-surgical OM directly assess the ability of vancomycin-saturated scaffolds to prevent infection in a contaminated bone defect [7, 8, 24]. Specifically, a 1 cm segment of bone was surgically excised from the middle of the right radius leaving the ulna intact. The segment was discarded and 1 × 10^6^ colony-forming units (CFU) of the *S. aureus* strain UAMS-1 [21] in 10 μL of PBS was injected into the medullary canal on each end of the exposed defect, thus resulting in a total inoculum of 2 × 10^6^ CFU. The surgical void created in the bone was then filled with scaffold saturated with PBS or PBS containing vancomycin. Control groups included uninfected rabbits exposed to scaffolds saturated with PBS or PBS saturated with 100 mg/mL of vancomycin. All groups in each experiment included 3 rabbits per group. Radiographic images were obtained on the surgical day (day 0) and at weekly or biweekly intervals thereafter.

At the completion of each experiment, rabbits were humanely euthanized and the surgical site exposed. To preserve the surgical site for μCT analysis, bacteriological sampling in these early experiments was limited to thoroughly swabbing the exposed surgical site and plating on tryptic soy agar (TSA) plates (Becton Dickinson, Franklin Lakes, NJ) without antibiotic selection. Plates were incubated overnight at 37°C. The right forelimb was then harvested and soft tissues removed before placing the bone in formalin until μCT analysis. In later experiments, the bone was stored at −80°C, scanned while frozen, and then pulverized for bacteriological analysis as detailed below.

### Microcomputed tomography (uCT)

After storage in formalin or at −80°C, image acquisition and analysis were done as previously described with minor modification [22, 26, 27]. Briefly, imaging was done via the Skyscan 1174 X-Ray Microtomograph (Bruker, Kontich, Belgium) with an isotropic voxel size of 11.6 μm, a voltage of 50 kV (800 μA) and a 0.25 mm aluminum filter. Samples that had not been stored in formalin were then subjected to bacteriological analysis. All μCT components were cleaned with 70% ethanol before and after use.

Reconstruction was performed with the Skyscan Nrecon software (Bruker, Kontich, Belgium). The reconstructed cross-sectional slices were processed with the Skyscan CT-analyzer software using the following steps: first, bone tissue was isolated from the soft tissue and background using a global thresholding (low = 85; high = 255). We then delineated regions of interest (ROI) by manually contouring the areas that contained scaffold. Next, we obtained the bone volume present in the drawn ROIs using the 3D analysis tool. In addition, we performed μCT on the S1 and K20 scaffolds prior to implantation (Fig 1) and calculated the bone volume, which allowed us to calculate the percent of scaffold remaining after implantation by comparison to the volume observed before implantation.

**Fig 1.**
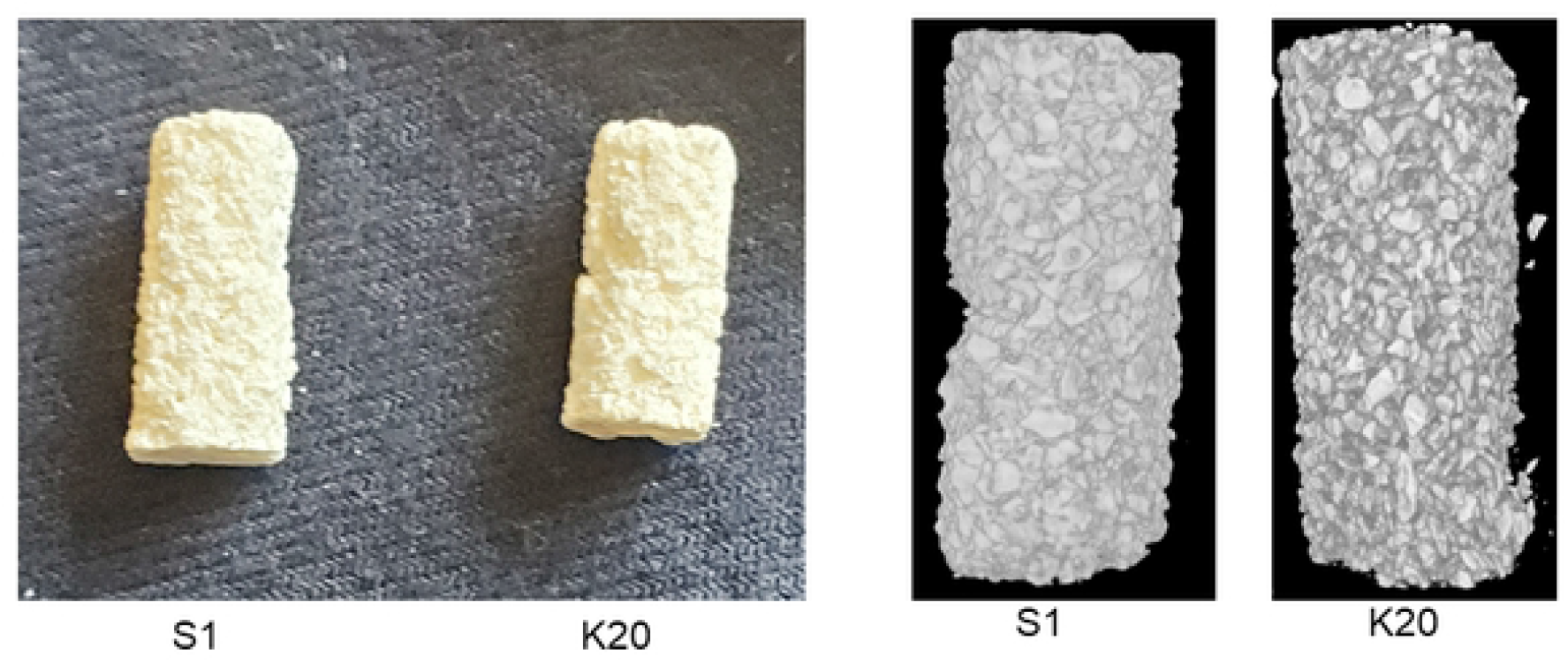
Schematic illustration of the polyurethane-based bone filler scaffold. Left: Photograph of the S1 and K20 scaffolds formulated in a size and shape to match the bone defect generated in our rabbit model. Right: μCT image of the S1 and K20 scaffolds prior to antibiotic loading.

### Assessment of bacterial burdens in bone

In short-term 14 day experiments in which μCT was not done and in the later longer term study in which μCT was done on frozen samples rather than formalin-fixed samples, dissected soft tissues including the remaining scaffold and the bone extending 0.5 cm on either side of the surgical site were homogenized using a Bullet Blender 5 Gold (Next Advance Inc., Troy NY). Aliquots of the homogenate were then plated on TSA plates to assess the number of viable bacteria. In all experiments, the identity of the *S. aureus* strain isolated at the end of the experiment was confirmed as the same strain used to initiate the infection by polymerase chain reaction (PCR) analysis to detect the *cna* gene, which encodes a collagen-binding surface protein that is not present in most strains of *S. aureus* but is present in the chromosome of UAMS-1 [24, 28, 29].

### Ethics statement

All animal experiments were approved by the Institutional Animal Care and Use Committee of the University of Arkansas for Medical Sciences (Animal Use Protocol 3896) and performed in compliance with NIH and FDA guidelines, the Animal Welfare Act, and United States federal law. During surgeries, buprenorphine was administered to minimize pain and the procedures were done under Xyalzine/Ketamine anesthesia maintained with inhaled isoflurane. Euthanasia was performed by a veterinarian using Euthasol. All efforts were made to minimize suffering.

### Statistical analysis

Statistical analysis of μCT results was done by unpaired t-test using GraphPad Prism 7.2 software (GraphPad Software Incorporation, La Jolla, CA, USA).

## Results and discussion

The dry weight of individual S1 and K20 scaffolds ranged from 84.78-90.61 and 84.62-90.27 mg, respectively. However, μCT imaging of each formulation before antibiotic loading provided visual evidence that the S1 scaffold was more dense than the K20 scaffold (Fig 1), and based on the amount of PBS remaining after saturation of each scaffold formulation the K20 scaffold was found to take up an average of approximately 2.30-fold more fluid than the S1 scaffold (range = 195-363 and 105-138 μL, respectively), thus confirming that the modifications made to the K20 scaffold increase its surface area and porosity relative to the S1 scaffold.

With both scaffold formulations, the amount of fluid uptake was consistent irrespective of antibiotic concentration, and no difference was observed in the concentration of antibiotic in the fluid remaining after saturation of the scaffold (data not shown). Thus, the increased amount of fluid uptake observed with the K20 scaffold was directly reflected in an increased amount of antibiotic uptake evident in both the greater amount of active vancomycin observed in the eluate after a 30 minute elution period (~20 vs ~10 mg/mL) and in the overall elution profile as assessed over time (Fig 2). For both the S1 and K20 scaffolds loaded at concentrations of 10 or 100 mg/ml of vancomycin, the concentrations antibiotic in the eluted samples obtained on day 10 remained above the breakpoint minimum inhibitory concentration (MIC) that defines a vancomycin-sensitive strain of *S. aureus* (≤2.0 μg/mL). However, the highest concentration (average ≥ 6.25 μg/ml) was observed with the K20 scaffold saturated with 100 mg/ml vancomycin, hereinafter designated as K20/100. The test strain in all cases, including the subsequent *in vivo* studies, was the methicillin-sensitive, USA-200 strain of *S. aureus* UAMS-1. This strain was chosen because it was obtained from the bone of a patient with OM during surgical debridement, forms a robust biofilm, has been proven to be virulent in relevant animal models of OM, and is clinically defined as sensitive to all of the antibiotics tested [8, 21–24]. We chose to focus on vancomycin for our *in vivo* studies because it is the primary antibiotic used to combat *S. aureus* infections caused by methicillin-resistant strains and thus would offer broader therapeutic utility.

**Fig 2.**
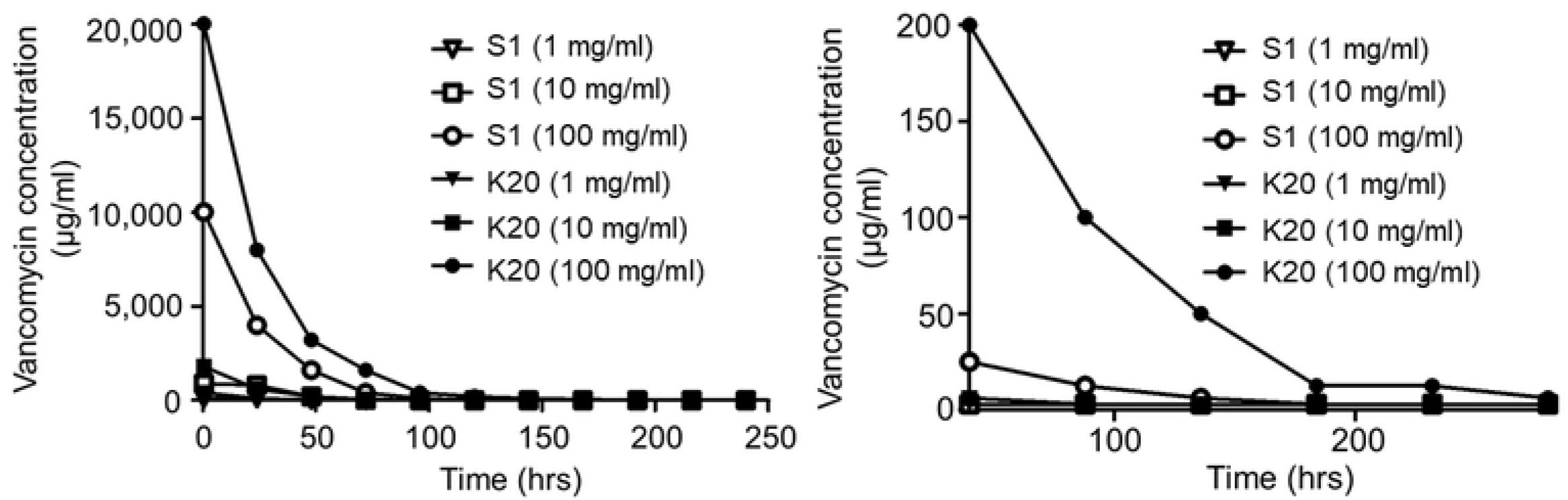
Uptake and elution of vancomycin as a function of scaffold formulation and antibiotic concentration. The S1 and K20 scaffold formulations were exposed to solutions of vancomycin at the indicated concentrations for 24 hours. Scaffolds were removed and placed in PBS without vancomycin. The buffer was replaced with antibiotic-free buffer at the indicated time points and the approximate amount of active vancomycin determined using a broth microdilution assay. Left: Results from the entire 10-day elution period. Right: exploded view of data focusing on the later time points.

To determine whether the amount of vancomycin observed with the K20/100 formulation could be defined as saturating, we also exposed the K20 scaffold to a solution containing 250 mg/mL of vancomycin, resulting in a greater initial release as defined after a 30 minute elution period (~32 mg/mL). However, within 24 hours the amount of antibiotic in the eluate fell to a concentration comparable to that observed with the K20 scaffold saturated with 100 mg/mL antibiotic (~8.0 mg/mL). Thus, as defined by the overall elution profile, we considered the K20 scaffold to be saturated when it was exposed for 24 hours to a solution containing 100 mg/mL of vancomycin. Although our *in vivo* studies were limited to vancomycin-saturated scaffold (see below), using this assay we also demonstrated that the K20 scaffold could be saturated with daptomycin (Fig S1) and gentamicin (Fig S2) in a concentration-dependent manner, although saturation required exposure to at least a 200 mg/mL solution for these antibiotics. These findings suggest that antibiotic-specific saturation parameters would need to be established prior to making recommendations regarding the clinical use of our bone regeneration scaffold as an anti-infective.

The antibiotic concentrations reported for these *in vitro* elution studies are defined by the nature of the experimental system, most notably the volume of buffer used for antibiotic elution, and so these results only allow relative conclusions about uptake and release as a function of scaffold formulation and antibiotic concentration. Thus, the critical question is whether these antibiotic concentrations are germane in the context of preventing infection in a contaminated bone defect. To address this, and to gain insight into whether the incorporation of vancomycin into the scaffold compromised its bone regenerative properties, we carried out three sequential *in vivo* experiments. All of these experiments employed the K20/100 scaffold, the difference being the length of time the experiment was extended after surgery and placement of the scaffold. This time period included 28, 56, and 84 days in the 1^st^, 2^nd^, and 3^rd^ experiments, respectively.

Given an experimental group size of 3 rabbits per group, these experiments collectively included 9 rabbits per experimental group. In the experimental group in which *S. aureus* was introduced and no antibiotic was added to the scaffold, the soft tissues surrounding the surgical site were confirmed to be heavily infected in 8 of 9 rabbits (Fig 3, Group 1). The exception was a single rabbit evaluated at 84 days post-infection, thus suggesting that the infection in this rabbit spontaneously resolved over time. These results confirm that the inoculum was sufficient to establish infection, particularly given that all isolates obtained at the end of each experiment were confirmed by PCR to be the same *S. aureus* strain used to initiate the infection (data not shown). In contrast, none of the 9 rabbits in the experimental group inoculated with *S. aureus* and treated with vancomycin-loaded scaffold were found to be infected at any time point (Fig 3, Group 2). Although bacteriological analysis in these studies was limited to swabs taken from the surgical site and did not include the bone or scaffold itself, these results nevertheless demonstrate that vancomycin can be incorporated into the K20 scaffold in an amount capable of limiting, if not preventing, infection. No bacteria were isolated from any of the 18 rabbits in either of two experimental groups that were not inoculated with *S. aureus* (Fig 3, Groups 3 and 4, respectively).

**Fig 3.**
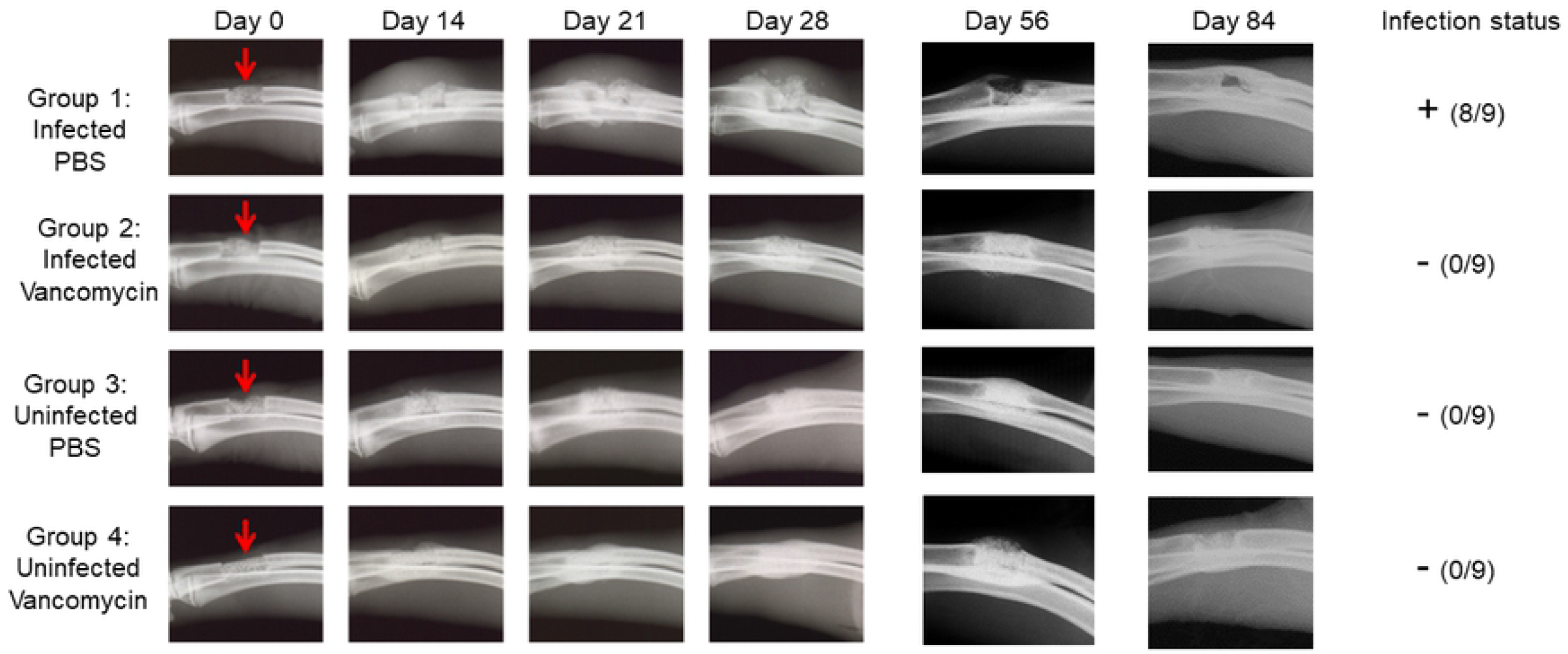
Bacteriological and radiographic evidence of osteomyelitis in experiments with the K20/100 scaffold formulation. A 1 cm defect (red arrow) was created in the right radius of rabbits as previously described [24]. The excised bone was discarded, and the distal and proximal ends of the remaining bone were either infected with *S. aureus* or inoculated with sterile PBS. The K20 formulation saturated with either PBS or vancomycin (100 mg/mL) was then placed into the defect and the wound closed. Left: Radiographs from the indicated time points are grouped by infection status and whether the scaffold was loaded with PBS or vancomycin. Three independent experiments were performed with end points at 28, 56 or 84 days. At each end point, rabbits were humanely euthanized, the wound opened, and swabs taken for bacteriological analysis. Right: The collective results of the bacteriological analysis.

Radiographic imaging from infected rabbits in which the surgically excised bone fragment was replaced with an anatomically-correct scaffold saturated with PBS also confirmed that the bacterial inoculum was sufficient to initiate an infection that progressed in severity in all three experiments, with radiographic evidence of infection being apparent as early as 14 days (Fig 3, Group 1). In contrast, there was little radiographic evidence of infection in any of the other experimental groups (Fig 3, Groups 2-4). Taken together with the results of our soft tissue bacteriological analysis, these findings suggest that the amount of antibiotic in the K20/100 scaffold formulation is sufficient to prevent infection from developing in a contaminated bone defect.

The μCT analysis confirmed that the scaffold was largely destroyed in infected rabbits that received PBS saturated scaffold (Fig 4, Group 1), while it remained in place and intact in all other experimental groups (Fig 4, Groups 2-4). These data suggest that the utility of the scaffold would in fact be compromised in an infected bone defect in the absence of antibiotic, and that the addition of vancomycin even in the saturating amount used in the K20/100 formulation did not adversely affect the bone regenerative properties of the K20 scaffold.

**Fig 4.**
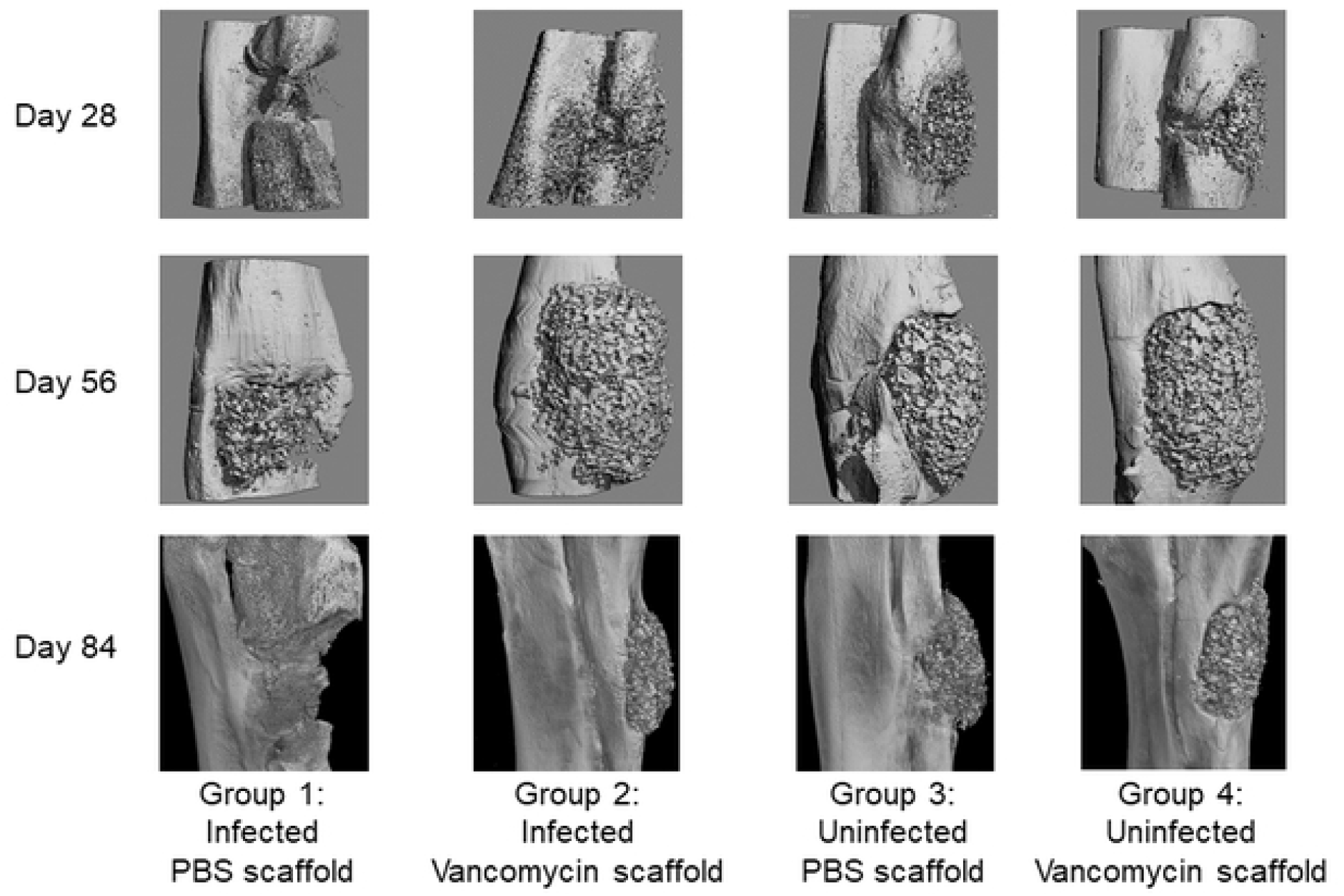
μCT analysis of K20/100 scaffold formulation *in situ*. Bone surrounding the surgical site was harvested 28, 56, or 84 days post-surgery and subjected to μCT analysis. Results shown are from representative rabbits from each experimental group, with the specific experimental group indicated below each set of vertical panels.

Quantitative analysis of bone regeneration was compromised in these experiments for two reasons. First, our rabbit model offers the advantage of allowing us to generate a segmental bone defect without disabling the rabbit because the radius and ulna are directly connected via an interosseous membrane that runs between the medial aspects of the bones (the radioulnar fibrous joint), thus providing structural stability even after creating a 1 cm segment defect in the radius. However, the presence of the intact ulna can also present a disadvantage in that it makes it difficult to accurately assess bone regeneration across the segmental defect itself. Second, the scaffold itself contains bone particles that are readily detectable by μCT (Fig 1). To compensate for these factors, we focused on the scaffold itself rather than the bone, and the results of these studies led us to conclude that approximately 50% of the scaffold remained after 84 days in both of the uninfected experimental groups (Fig 5, Groups 3 and 4). The amount of scaffold remaining in infected rabbits exposed to vancomycin-loaded scaffold was slightly higher (Fig 5, Group 2), which suggests that the transient presence of bacteria may have slowed decomposition of the scaffold. However, the difference did not reach statistical significance by comparison to the uninfected experimental groups. As suggested by visual analysis of μCT images (Fig 4), the analysis also confirmed that the scaffold was essentially destroyed in infected rabbits exposed to PBS-saturated scaffold (P = 0.0274) (Fig 5, Group 1).

**Fig 5.**
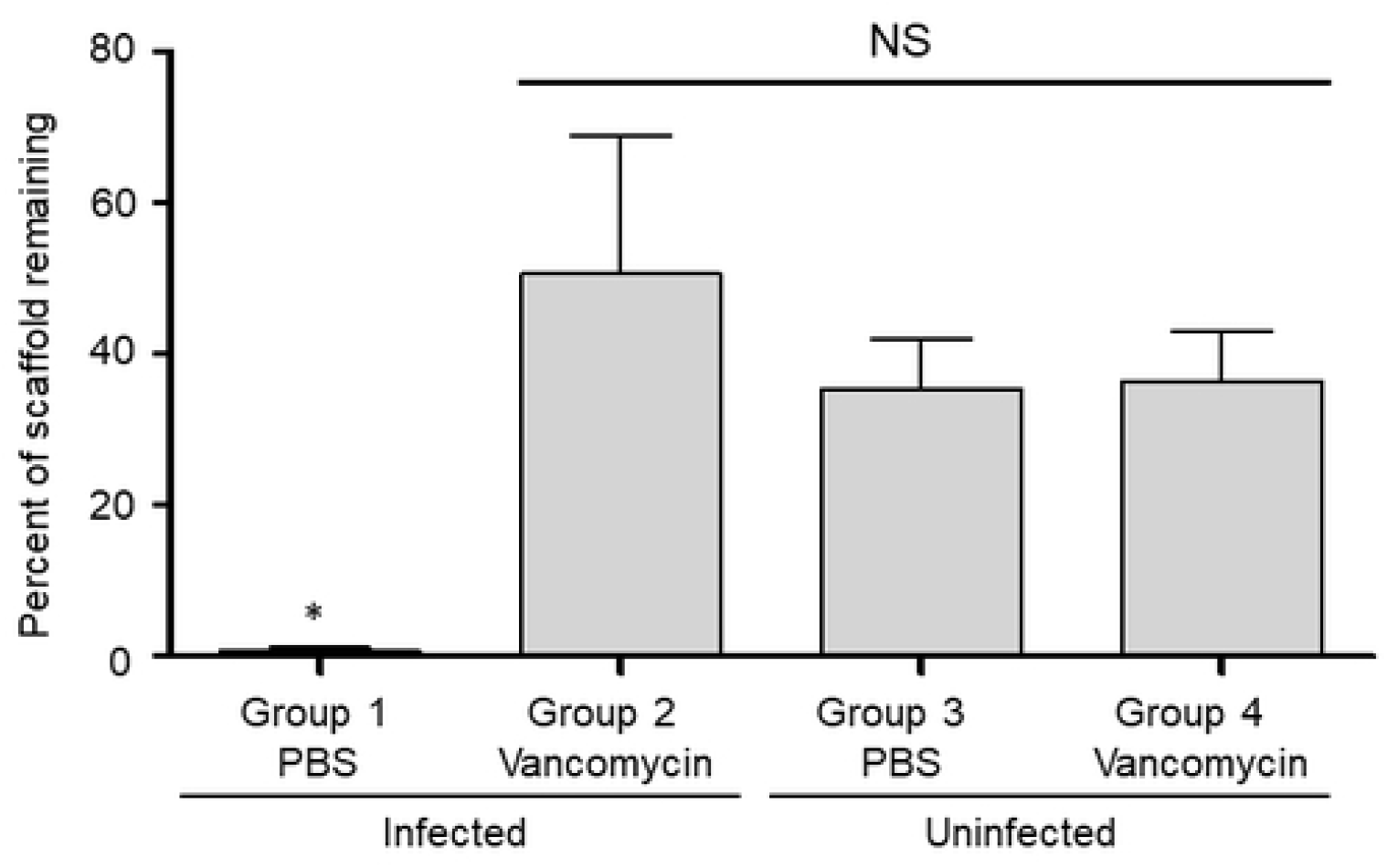
Percentage of K20/100 scaffold formulation remaining as a function of infection status and antibiotic loading. Bone surrounding the surgical site was harvested after 84 days and subjected to μCT analysis. The amount of scaffold remaining was calculated by comparison to control scaffolds prior to implantation (Fig 1). Results shown are the average ± the standard error of the mean for each experimental group. Asterisk indicates a statistically significant difference (P= 0.0274) by comparison to the uninfected control group in which the surgically-created bone defect was not contaminated and filled with PBS-saturated scaffold (Group 3).

Statistical analysis was limited by the small size of each experimental group in each of the three sequential experiments and the inability to combine datasets given the use of different time frames used in each experiment. However, the overall results of all three experiments are consistent with each other. Specifically, in the experimental group in which rabbits were infected with UAMS-1 and exposed to PBS-saturated scaffold (Fig 5, Group 1), the same bacterial strain used to initiate the infection was isolated from 8 of 9 rabbits at the end of each experiment, there was clear evidence of OM as assessed by radiographic analysis (Fig 3), and the segmental defect was not filled as assessed by μCT (Fig 4). In contrast, no bacteria were isolated from any of the 27 rabbits in the other three experimental groups, there was no radiographic evidence of the development of OM in any of these rabbits (Fig 3), and there was no significant difference in bone regeneration in rabbits in any of these three groups (Fig 4 and Fig 5). Thus, the results of these three independent experiments are all consistent with the hypothesis that it is possible to incorporate vancomycin into the K20 scaffold in an amount sufficient to prevent infection from developing in a heavily contaminated segmental bone defect without compromising the bone regenerative properties of the scaffold itself.

The bacteriological analysis of these experiments was restricted to soft tissue swabs of the surgical site, thus leaving open the possibility that viable intraosseous bacteria were not eradicated. Additionally, the observation that the vancomycin-saturated K20/100 formulation is sufficient to prevent infection from developing in a contaminated bone defect does not mean that the amount of vancomycin used in these studies is necessary to do so, and there are important reasons to consider the use of less antibiotic. Moreover, the use of less antibiotic would minimize the possibility that localized antibiotic delivery would contribute to systemic toxicity, particularly given the clinical reality that localized antibiotic delivery is rarely if ever used without concomitant systemic antibiotic therapy. However, given the tremendously adverse consequences of infection, the use of less antibiotic is dependent on the knowledge that less antibiotic is sufficient to reliably clear a contaminated bone defect.

To address these issues, we carried out two experiments with three significant modifications that allowed us to focus specifically on determining the minimum amount of vancomycin required to prevent infection. First, infected rabbits were treated using the K20 scaffold saturated with PBS containing 100, 75, 50, 25 and 0 mg/mL of vancomycin. Second, by comparison to the loading protocol described for the 1^st^ three experiments, scaffolds were soaked in the vancomycin solutions for 1 rather than 24 hours. This reduction in loading time was based on our demonstration that fluid uptake in the K20 scaffold is virtually immediate and the logic that a shorter loading period would facilitate the intraoperative use of the scaffold. However, the time reduction was done only after confirming that 1 hour was sufficient to achieve an elution profile directly comparable to that observed after 24 hours (data not shown). Because we were evaluating scaffolds loaded using different concentrations of vancomycin, in addition to our bioassay, we also used a spectrophotometric assay [25] as a means of assessing the absolute concentration of vancomycin released from the K20 scaffold. Studies done with this assay confirmed a linear relationship between vancomycin concentration and absorbance at vancomycin concentrations between 0.4 and 50 mg/mL (Fig S3). For those samples in which our bioassay indicated a concentration higher than 50 mg/mL, samples were diluted to get within the linear range. The absorbance was then used to calculate the amount of vancomycin in the original sample.

The results of these spectrophotometric assays were generally consistent with the results of our bioassays, although when viewed collectively there was evidence to suggest that our bioassays may have underestimated the amount of antibiotic (Fig S4). This finding is not surprising given our use of two-fold dilutions in our bioassays and the fact that we chose to report the lower concentration of the range defined by these dilutions. Analysis also confirmed that antibiotic uptake and release is dependent on the concentration of antibiotic in the solution used to saturate the scaffold (Fig S4). These results demonstrate that antibiotic loading can be controlled by modifications to the scaffold itself (Fig 2) and by altering the concentration of antibiotic used to load the scaffold (Fig S4), thus providing further flexibility in the use of our scaffold as an anti-infective.

Finally, because the focus in these experiments was solely on preventing infection, the experiments were terminated 14 days after surgery and bacteriological analysis was extended to include the surgical site and the bone itself. Because any implanted material containing antibiotic can become a substrate for bacterial colonization as the level of antibiotic is diminished, we also included the scaffold itself in our bacteriological analysis. The use of the scaffold and bone in the bacteriological assay precluded μCT analysis and, given the time frame for the development of radiographic evidence of OM, limited the information that could be gained by X-ray. However, use of the complete area did allow for exhaustive bacteriological analysis. As in our earlier experiments, all six rabbits in the untreated control group were culture positive based on swabs of the exposed surgical site, while no bacteria were isolated from any rabbits in any of the other experimental groups (Fig 6). The bone in all six rabbits in the infected but untreated experimental group was also found to be heavily colonized. Viable bacteria were also isolated from the bone and/or scaffold of 7 of 24 of the infected rabbits treated with scaffold loaded with varying concentrations of vancomycin. Of these 7, four were treated with the K20/25 scaffold, with the K20/50, K20/75, and K20/100 experimental groups containing 0, 1, and 2 culture-positive rabbits, respectively (Fig 6). However, the presence of even a single colony was counted as a positive, and in all 7 cases the bacterial load was dramatically reduced by comparison to the number isolated from all 6 rabbits in the infected, untreated experimental group (Fig 6). This reduction, along with the clearance seen with soft tissue samples in the previous experiments (Fig. 3) suggests that clearance may have been possible if the experiment had be continued past 14 days.

**Fig 6.**
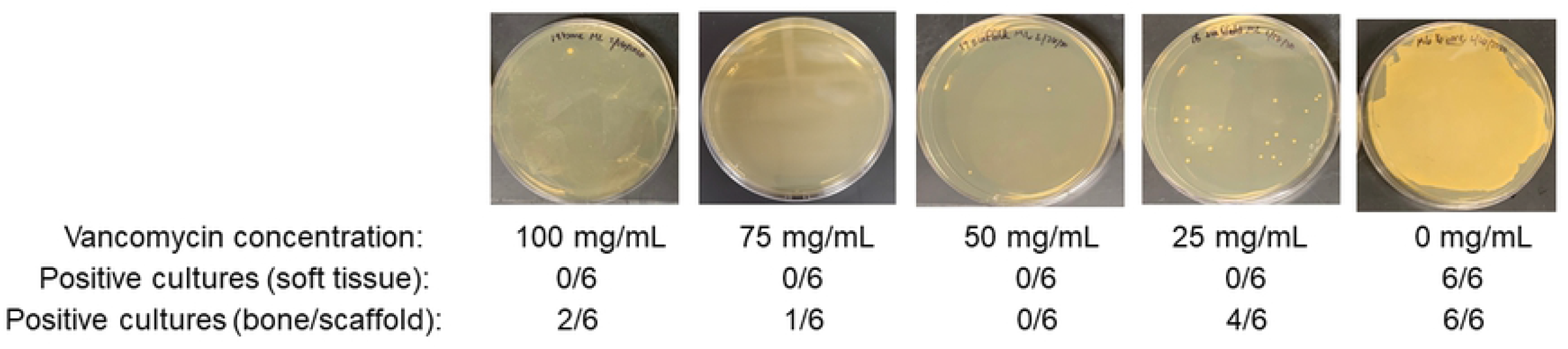
Bacteriological analysis of the K20 scaffold as a function of antibiotic concentration. The K20 scaffold was saturated in a solution containing the indicated amounts of vancomycin for 1 hour at room temperature. Scaffolds were then used to replace a 1 cm excised bone segment in infected rabbits. Scaffolds were left in place for 14 days, at which point the surgical site was opened and soft tissues samples swabbed for bacteriological analysis. The bone and its associates scaffold was then removed and homogenized before plating an aliquot on tryptic soy agar. All soft tissue samples were negative as indicated below each panel. Images shown for each antibiotic concentration are representative of rabbits in each experimental group.

These collective studies suggest that loading the K20 scaffold with 100 mg/ml of vancomycin does not compromise its bone regenerative properties in comparison to scaffolds loaded with PBS in the absence of infection. Our findings also suggest that a significant anti-infective benefit can be achieved with the K20 scaffold loaded with only 50 mg/mL of vancomycin. In our initial antibiotic loading studies (Fig 2), the S1 scaffold was found to have a loading capacity ~50% of that observed with the K20 scaffold. The prevention of infection by K20 at half the potential loading capacity suggests that the S1 scaffold saturated with vancomycin would also be sufficient to minimize infection in a contaminated bone defect. To test this idea, we carried out a final experiment that differed from previous experiments in two respects. First, we used the S1 scaffold saturated in 100 mg/mL of vancomycin rather than the K20/100 formulation. Second, radiographs were taken every 21 days throughout the 84 day post-infection period, at which point we employed a modified protocol utilizing frozen rather than formalin-fixed bones in an attempt to obtain μCT and comprehensive bacteriological data from the same rabbit.

In this study, the only experimental group in which viable bacteria were isolated from soft tissue swabs of the exposed surgical site were in the infected experimental group in which the surgically-removed defect was replaced with PBS-saturated scaffold, with all three rabbits in this group being heavily culture positive (Fig 7, Group 1). Based on bacteriological analysis of the bone/scaffold, we did confirm one culture positive rabbit in the S1/100 experimental group, but the number of viable bacteria obtained from this rabbit was <10^2^, while in the corresponding experimental group treated with the PBS-saturated scaffold the number ranged from 10^4^ to 3 × 10^7^ (data not shown). These results are consistent with radiographic analysis, which revealed clear signs of soft tissue infection and osteonecrosis in all rabbits in this experimental group (Fig 7, Group 1) and the absence of these signs in rabbits from all other experimental groups (Fig 7, Groups 2-4). Radiographic analysis also confirmed that the bone defect remained unfilled in the Group 1 rabbits but that this was not the case in rabbits in any other experimental group (Fig 7). As with the K20 scaffold, the amount of the S1 scaffold remaining after 84 days was also significantly lower (P = 0.0375) in all Group 1 rabbits by comparison to all other experimental groups, and comparable among all other experimental groups irrespective of the presence or absence of vancomycin (Fig 8). However, as with the K20/100 scaffold, there was a trend suggesting that the presence of bacteria may have delayed incorporation of the scaffold, although once again this trend did not reach statistical significance (Fig 8).

**Fig 7.**
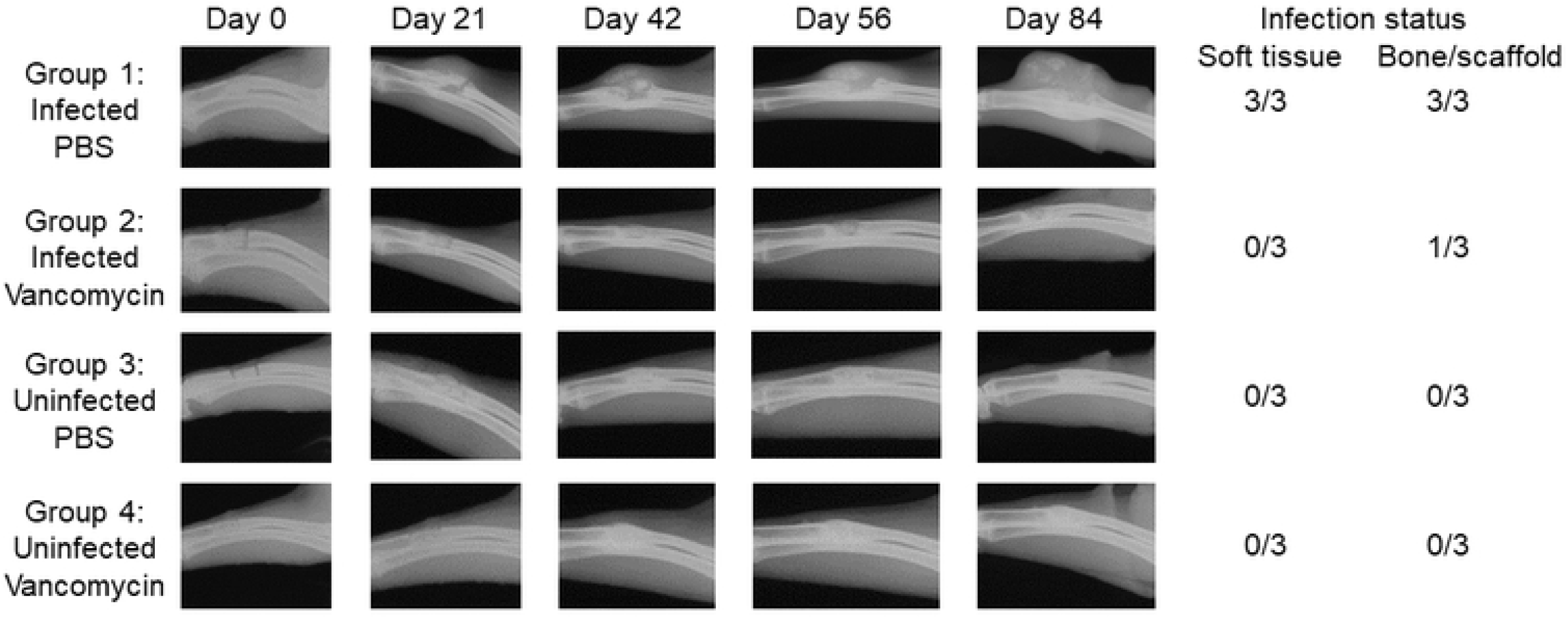
Radiographs from representative rabbits in each experimental group treated with the S1/100 scaffold formulation. Representative X-rays are shown from infected and uninfected rabbits exposed to the S1 scaffold saturated with PBS or 100 mg/mL vancomycin as indicated to the left. After 84 days, rabbits were humanely euthanized and the surgical limb harvested and frozen. Upon removal from the freezer, the surgical site was exposed and swabs taken from the surrounding soft tissues. The bone and its associated scaffold were then imaged by μCT before being homogenized and plated on tryptic soy agar to recover viable bacteria. The collective results observed with soft tissue samples and the bone/scaffold are shown to the right.

**Fig 8.**
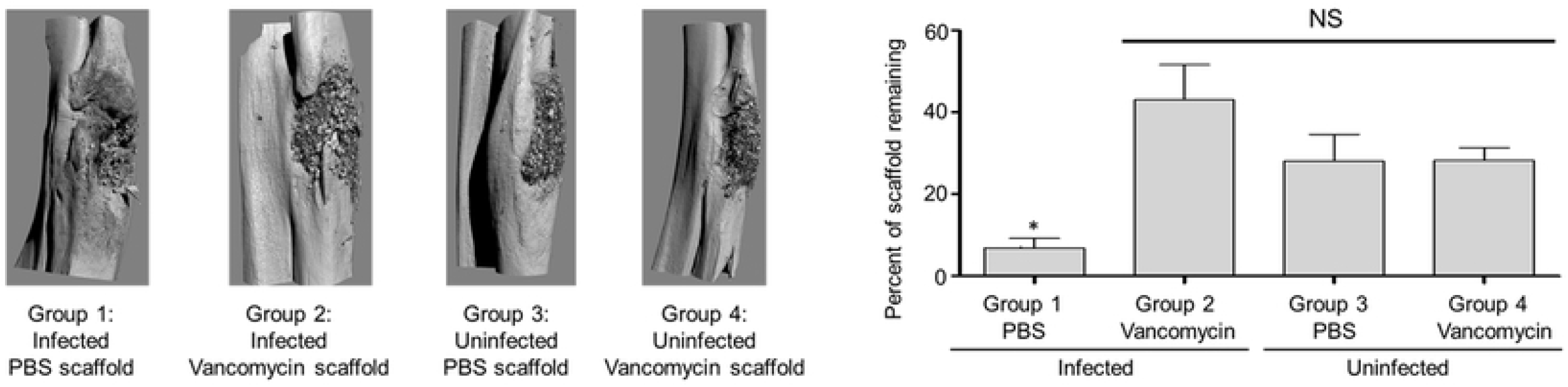
μCT analysis of S1/100 scaffold *in situ*. Left: Bone surrounding the surgical site was harvested and subjected to μCT analysis. Results shown are from representative rabbits from each experimental group, with the specific experimental group indicated below each panel. Analysis is based on results observed after 84 days. Right: The amount of scaffold remaining at the surgical site was calculated by comparison to control scaffolds prior to implantation. Results shown are the average ± the standard error of the mean for each experimental group. Asterisk indicates a statistically significant difference (P = 0.0375) by comparison to the uninfected control group that obtained PBS-saturated scaffold (Group 3).

In summary, the studies we report here were carried out as six independent experiments that differed with respect to the scaffold formulation used (K20 vs. S1), the time after surgery at which the results were evaluated (14, 28, 56 and 84 days), the amount of vancomycin used to saturate the scaffold (25-100 mg/ml), and the method employed for bacteriological analysis (soft tissue vs. soft tissue and bone/scaffold). These changes were implemented to address specific issues as they arose as our experiments progressed. While necessary, the changes between experiments did compromise statistical analysis owing to the relatively small group size in each experiment and the inability to combine the results of different experiments. However, the collective results strongly support the hypothesis that antibiotics, specifically vancomycin, can be incorporated into our polyurethane-based bone filler scaffold at levels that prevent or at least significantly limit the development of infection even in a heavily contaminated bone defect without compromising the bone regenerative properties of the scaffold.

That said, we acknowledge the limitations of these studies. We did not continue the experiments post 84 days to fully assess the bone formation in every case. In addition, we did not observe complete resorption of the scaffolds even in the experiments extended to 84 days, although significant resorption was evident. However, we have seen excellent regenerative properties of the scaffolds, as demonstrated in other models, [15–18] and we believe the collective studies we report do allow us to conclude that the inclusion of vancomycin even at saturating levels did not compromise these properties. Our data does suggest that the presence of bacteria may delay replacement of the scaffold with bone, but this delay was not statistically significant, and more importantly must be interpreted in the context of the extremely adverse consequences associated with the failure to prevent infection as evidenced by the results observed in infected rabbits exposed to the scaffold without antibiotic. We would also emphasize that the scaffold formulations we examined provide a starting point to carry out the proof-of-principle studies we report, and therefore are an important and necessary experimental foundation to move forward, although they certainly do not preclude the possibility of additional modifications to the scaffold formulation that would further enhance both its antibiotic delivery and bone regenerative properties.

We were also unable to demonstrate that we could completely eradicate all viable bacteria from all experimental animals, particularly at 14 days post implantation and with lower drug concentrations. However, even using highly stringent experimental methods that allowed us to quantitatively isolate bacteria from the bone, the surrounding soft tissues, and the scaffold itself, we did demonstrate that the saturation of the K20 or S1 scaffolds with ≥50 mg/ml of vancomycin eliminated viable bacteria from most of the experimental animals (20 of 24, 83.3%) and, even in rabbits in which this was not the case, the number of viable bacteria recovered from each rabbit was reduced by several orders of magnitude by comparison to untreated controls. Additionally, there was no radiographic evidence of OM even in the few culture-positive rabbits observed in the infected experimental groups treated with vancomycin-saturated scaffold. This finding is particularly true given that we employed an experimental protocol in which the bone defect was heavily contaminated with 2 × 10^6^ viable bacteria introduced directly into the intramedullary canal, thus allowing us to ensure that the infection was established in the absence of treatment and to stringently assess the ability to limit if not completely prevent this extremely adverse consequence of traumatic injuries to the bone. Thus, we believe the collective results we report are highly supportive of the conclusion that this polymeric-based bone filler scaffold can be used as an effective antibiotic delivery matrix without significantly compromising its bone regenerative properties. The importance of this conclusion is emphasized by the need for alternative bone fillers that can be produced in a stable, readily available form and rapidly loaded with the antibiotic of choice, thereby providing orthopaedic and trauma surgeons with a means of both preventing the extremely adverse consequence of infection following traumatic injury to the bone and while simultaneously providing a means of promoting bone repair.

## ACKNOWLEDGEMENTS

This work was supported by grants from the Arkansas Research Alliance and a generous gift from the Texas Hip and Knee Foundation.

## SUPPORTING INFORMATION FIGURE LEGENDS

**Fig. S1. Uptake and elution of daptomycin as a function of antibiotic concentration.** The K20 scaffold was exposed to solutions of daptomycin at the indicated concentrations for 24 hours. Scaffolds were then removed and placed in PBS without antibiotic. Scaffolds were then removed and placed in PBS without daptomycin. The buffer was replaced with fresh antibiotic-free buffer at the indicated time points and the approximate amount of active daptomycin determined using a broth microdilution bioassay. Left: the results from a 7-day elution period. Right: Exploded view of the data focusing only on the later time points.

**Fig. S2. Uptake and elution of gentamicin as a function of antibiotic concentration.** The K20 scaffold was exposed to solutions of gentamicin at the indicated concentrations for 24 hours. Scaffolds were then removed and placed in PBS without antibiotic. Scaffolds were then removed and placed in PBS without gentamicin. The buffer was replaced with fresh antibiotic-free buffer at the indicated time points and the approximate amount of active gentamicin determined using a broth microdilution bioassay. Left: the results from a 7-day elution period. Right: exploded view of the data focusing only on the later time points.

**Fig S3. Correlation between concentration of active vancomycin and absorbance at 280 nm.** A solution of vancomycin was freshly prepared in PBS at the indicated concentrations and the absorbance at 280 nm measuring as a function of antibiotic concentration.

**Fig. S4. Correlation between bioassay and spectrophotometric assay for vancomycin.** The K20 scaffold formulation was saturated in a solution containing the indicated amounts of vancomycin for 1 hour at room temperature. Scaffolds were then removed and placed in PBS without vancomycin. The buffer was replaced with fresh antibiotic-free buffer at the indicated time points and the approximate amount of active vancomycin at each sampling point determined by bioassay (MIC) and spectrophotometric analysis (Spec).

